# Multimodal Magnetic Resonance Histology and Light Sheet Imaging for Quantitative Neurogenetics of the Mouse

**DOI:** 10.1101/2021.10.08.463660

**Authors:** G. Allan Johnson, Gary Cofer, James Cook, James Gee, Adam Hall, Kathryn Hornburg, Yi Qi, Yuqi Tian, Fang-Cheng Yeh, Nian Wang, Leonard White, Robert W. Williams

**Affiliations:** Duke University; University of Pennsylvania; Life Canvas Technologies Inc; University of Pittsburgh; Indiana University School of Medicine; University of Tennessesse

## Abstract

Paul Lauterbur closed his seminal paper on MRI with the statement that “zeugmatographic (imaging) techniques should find many useful applications in studies of the internal structures, states and composition of microscopic objects” [1]. Magnetic resonance microscopy was subsequently demonstrated in 1986 by three groups [2] [3] [4]. The application of MRI to the study of tissue structure, i.e. magnetic resonance histology (MRH) was suggested in 1993 [5]. MRH, while based on the same physical principals as MRI is something fundamentally different than the clinical exams which are typically limited to voxel dimensions of ~ 1 mm^3^. Preclinical imaging systems can acquire images with voxels ~ 1000 times smaller. The MR histology images presented here have been acquired at yet another factor of 1000 increase in spatial resolution. Figure S1 in the supplement shows a comparison of a state-of-the-art fractional anisotropy images of a C57 mouse brain in vivo @ 150 μm resolution (voxel volume of 3.3 ×10^−3^ mm^3^) with the atlas we have generated for this work at 15 μm spatial resolution (voxel volume of 3.3 × 10^−6^ mm^3^). In previous work, we have demonstrated the utility of MR histology in neurogenetics at spatial/angular resolution of 45 μm /46 angles [6]. At this spatial/angular resolution it is possible to map whole brain connectivity with high correspondence to retroviral tracers [7]. But the MRH derived connectomes can be derived in less than a day where the retroviral tracer studies require months/years [8]. The resolution index (angular samples/voxel volume) for this previous work was >500,000 [9]. Figure S2 shows a comparison between that previous work and the new atlas presented in this paper with a resolution index of 32 million.

Light sheet microscopy (LSM) has undergone similar rapid evolution over the last 20 years. The invention of tissue clearing, advances in immunohistochemistry and development of selective plane illumination microscopy (SPIM) now make it possible to acquire whole mouse brain images at submicron spatial resolution with a vast array of cell specific markers [10] [11] [12] [13]. And these advantages can be realized in scan times of < 6hrs. The major limitation from these studies is the distortion in the tissue from dissection from the cranium, swelling from clearing and staining, and tissue damage from handling.

We report here the merger of these two methods:

1. MRH with the brain in the skull to provide accurate geometry, cytoarchitectural measures using scalar imaging metrics and whole brain connectivity at 15 μm isotropic spatial resolution with super resolution track density images @ 5 μm isotropic resolution;
2. whole brain multichannel LSM @ 1.8×1.8×4.0 μm;
3. a big image data infrastructure that enables label mapping from the atlas to the MR image, geometric correction to the light sheet data, label mapping to the light sheet volumes and quantitative extraction of regional cell density. These methods make it possible to generate a comprehensive collection of image derived phenotypes (IDP) of *cells and circuits covering the whole mouse brain* with throughput that can be scaled for quantitative neurogenetics.

## Results

All animal studies were approved by the Duke University Institutional Care and Use Committee. Figure 1 provides an overview of the workflow: tissue preparation, image acquisition, post processing, and data integration into the common space. There are two processing streams. The first focuses on MR histology with the brain in the skull. When the MRH data has been acquired, the brain is extracted from the skull and transferred to the second acquisition stream for light sheet microscopy. The two streams are merged into a common space to allow generation of image derived phenotypes of cells and circuits covering the whole brain.

### Computer infrastructure

A high-performance computer infrastructure has been assembled to facilitate the several pipelines described below. Figure S3 summarizes the elements of that infrastructure. Specific attention has been paid to the ability to handle multiple 3D arrays as large as 500 GB. Data streaming to a high-performance cluster (18 node, 604 processors) automates the reconstruction. Two additional high-performance servers with 1.5TB of memory allow interactive analysis of (4D) i.e. layered 3D data sets using multiple packages (Fiji, Slicer, Imaris). High performance RAID devices placed at crucial point in the acquisition/analysis chain further facilitate this interactive analysis.

### MR Histology Acquisition

MR histology (MRH) provides the key element of the method by providing an accurate geometric map of *each* specimen, yielding a common space into which all the data can be integrated. The brain is perfusion fixed using ProHance, (Gadoteridol), a chelated Gd compound that reduces the tissue spin lattice relaxation time from ~ 1800 ms to 100 ms. The reduced T1 allows use of shorter repetition intervals (TR) required for the microscopic resolution. The shorter TR combined with compressed sensing (described below) and long scan times have allowed us to acquire quantitative DTI data at 15 μm isotropic resolution and track density images at 5 μm which we believe to be the highest spatial resolution yet achieved.

MR images are acquired on a 9.4T/89mm vertical bore magnet controlled by an Agilent Direct Drive console (Vnmrj 4.0). Key to the success is a gradient insert constructed by Resonance Research Inc (Billerica, Ma). The BRG-88_41 coil provides peak gradients up to 2500mT/m with rise times of 25000T/m/s and ±5% linearity over a 30 mm diameter spherical volume. The high-capacity cooling system (600 W power dissipation) allows one to use exceptionally strong gradients (1500mT/m) with high duty cycle (TR=100 ms). The perfusion fixed brain is placed in a 10 mm diameter tube with a 3D printed insert to stabilize it in the container. The tissue is surrounded by fomblin, a fluorinated lubricant that minimizes the susceptibility artifacts at the surface of the brain. The container is placed in a Stesjkalrf coil constructed from a single sheet of silver which yields high Q (>600), high homogeneity, and high sensitivity. Two imaging sequences are used; 1) a multigradient echo (MGRE) sequence (TR/TE = 100/4.4ms) and a Stesjkal/Tanner spin echo sequence for diffusion tensor imaging (DTI) (TR/TE =100/12-19 ms) [14]. Both sequences employ phase encoding along the short axes of the specimen (x and y) with the readout gradient applied along the long axis of the specimen (z).

Compressed sensing has been employed to accelerate both sequences [15] [16]. A probabilistic map is generated for sparse sampling of Fourier space along the two-phase encoding axes (Figure 2). A script on the scanner starts the acquisition of the first 3D volume, a baseline image (b0) with no diffusion encoding gradient. At the conclusion of the scan, the data is automatically written to the Dell cluster. A Fourier transform is applied along the z axis (the long axis of the brain) producing between 256-3000 individual 2D files which are then queued for iterative reconstruction in the multiple processors on the cluster. When all the 2D files have been reconstructed, the data is reassembled into a 3D volume. While this is under way, the scanner launches the next volume in the sequence with a diffusion encoding direction (bvec) and value (bval) determined from a file accessed via the scanner script. The process proceeds under control of the scanner script through as many bvecs and bvals as one chooses. Baseline acquisitions are included in every 10-15 volumes. Twelve data sets were acquired with spatial resolution ranging from 15-25 μm, angular sampling ranging from 61 to 126 and b values of 3000 s/mm^2^. The acquisition protocols are summarized in Table 1.

### Registration-4D volume creation

The strong gradients used for diffusion encoding induce eddy currents in the magnet bore that spatially shift the individual 3D volumes. The shifts are dependent on the magnitude and direction of the diffusion gradients. A skull stripping algorithm is applied to each volume as a preprocessing step that facilitates registration of the multiple 3D volumes into a single 4D array. The next step registers all the baseline (b0) volumes together. The pipeline based on ANTs uses an affine transform with scaling and shearing to sequentially registers each 3D diffusion weighted volume to the baseline template [17] [18]. A second pass of the pipeline registers each diffusion weighted volume to the average baseline template.

### Denoising

The tradeoffs between signal to noise ratio, spatial resolution (voxel volume), diffusion weighting (b value), and scan time for a single 3D volume are known. Addition of multiple volumes i.e. angular sampling adds a fifth dimension. St-Jean et al have developed a novel denoising algorithm that recognizes the 4^th^ (angular) dimension of the data [19]. The volume is decomposed into 4D overlapping patches that sample both the spatial and angular resolution. A dictionary of atoms is learned on the patches and a sparse decomposition is generated by bounding the reconstruction error with the local noise variance. The method improves the visibility of structures while simultaneously reducing the number of spurious tracts. St-Jean and colleagues implemented the algorithm for high resolution clinical scans (matrix of 210×210×210) of ~ 20 MB/volume with 40 volumes (~800MB). The atlases generated for this work were acquired with arrays as large 511×711×1132 arrays (~900 MB/volume) with 108 angular samples plus 13 baseline images i.e. a 4D volume (~100 GB) that is more than 100 times larger. To accommodate the change of scale, the algorithm was implemented on the cluster (Figure S3) by breaking the volume into overlapping cubes which could be processed in parallel. Figure S4 shows the diffusion weighted images from specimen 200316-1:1 (Table 1) A) before and B) after denoising.

### Diffusion tensor calculation

The 4D volume is passed to DSI Studio (http://dsi-studio.labsolver.org/) where a perl script executes the initial pass using the diffusion tensor algorithm to generate five different 3D scalar volumes: axial diffusivity (AD), radial diffusivity (RD), mean diffusivity (MD), fractional anisotropy (FA), and color fractional anisotropy (clrFA) images. A MATLAB script averages all the diffusion weighted 3D volumes to produce the diffusion weighted image (DWI). The 4 echoes of the MGRE image are averaged together providing an image in which tissue contrast is driven by the transverse relaxation time and local field inhomogeneity (T2*-weighted). These seven 3D scalar images have decidedly different contrast highlighting different anatomical landmarks. The combination of MGRE and DTI scalar data drive the registration of the atlas and associated labels.

### 15 μm DTI Atlas

An atlas was created for both male (specimen ID: 200302-1:1, Table 1) and female (specimen ID: 200316-1:1 Table 1) 90-day C57BL/6J mouse. Figure 3 shows representative cross sections for the average MGRE, DWI, FA, and colorFA images of the female specimen with magnified regions of the hippocampus. Small vessels are evident in the MGRE and DWI. The dentate gyrus is most dramatically highlighted in the DWI along with the hippocampal fissure in a fashion like that seen on a Nissl section. The FA more closely resembles an acetylcholinesterase section. And the directional information from the diffusion eigenvectors in the color FA helps differentiate the boundary between CA2 and CA3.

The label set from the Allen Brain Atlas (ABA) common coordinate framework (CCF3) [20] has been mapped to both the male and female 15 μm MR data sets. As a first step, the entire collection of 461 labels (in each hemisphere) was registered to the MR volumes in ANTs using affine and diffeomorphic transforms. Many of the structures are so small that one would have difficulty reliably registering them across the diverse strains. Thus, a reduced set of labels (RCCF3) was created by combining smaller sub volumes resulting in a collection of 180 regions of interest for each hemisphere. The complete label set and relationship to CCF3 is included in the Supplement table S5. Three experienced neuroanatomists reviewed the delineations which were mapped onto six different scalar volumes (MGRE, DWI, AD, RD, FA, clrFA), all of which were registered to each other. The volume parcellation was refined by a) viewing the labels and underlying gray scale or color images simultaneously in the three cardinal planes; and b) enabling the editor to easily toggle between any of the registered volumes. A boundary that is poorly visible in the MGRE might be more readily visible in the DWI. Delineations were iteratively edited to be a) consistent across the collection of six scalar images; b) consistent in all three cardinal planes; consistent with the ABA reference atlas (Supplement Figure S5)

### Atlas Registration

The atlas and associated labels are registered to the strain under study using our previously published SAMBA pipeline built on the Advanced Normalization Tool set (ANTs). An initial rigid transformation aligns the unknown to the RCCF atlas described above. In population studies with multiple specimens of a given strain, a minimum deformation template (MDT) is formed through iterative affine and diffeomorphic transforms using multiple combinations of the scalar images. A combination of DWI and FA has proven effective for the studies we have undertaken to date. Once an MDT is complete, all specimens in that cohort are reregistered to it. Registration between images of similar contrast (FA to FA) are executed using the cross-correlation similarity metric. Registration between images of differing contrast (FA to DWI) use mutual information. When the atlas and labels have been mapped to the specimen under study, the transform is inverted to map the labels back to the specimen under study leaving a 4D volume with labels for the new specimen from which scalar image phenotypes can be derived. A MATLAB script extracts volume, and mean and standard deviation of the scalar diffusion metrics (AD, RD,MD,FA) for all 180 (x2) regions of interest. The pipeline calculates a coefficient of variation between left and right hemispheres as a quality assurance check. The summary of regional scalar image derived phenotypes is written to a summary spread sheet with the meta data for the study.

### Tractography

The 4D volume with labels, b vectors and b values are passed to DSI studio to generate whole brain tractography, track density images, connectome and associated statistics using the GQI algorithm [21]. The GQI algorithm exploits the higher angular sampling resolving multiple fibers (up to 4) in each voxel. Whole brain tractography is executed on CTX01 (Figure S3), a server with 1.5TB of memory, sufficient for the .fib file generated in DSI Studio.

### Light Sheet Acquisition/Transfer

The brain is extracted from the skull and placed in normal saline and shipped to Life Canvas Technology in an airtight container which minimizes the formation of bubbles in the brain during air shipping. The tissue is cleared using SHIELD [11] and stained using SWITCH [12]. The smart SPIM light sheet fluorescent microscope provides whole brain coverage (3650μm field of view) @ 1.8×1.8×4.0 μm with a 3.6 X objective producing three collections of registered 2 D .tiff files, one collection for each excitation wavelength. The three collections are transferred to Duke via Globus directly into the 8TB SSD array on the server configured for light sheet imaging.

### Light Sheet Registration

A dedicated pipeline on the LSM server registers the LSM data to the companion MRH volume. The first column in Figure 4 shows a raw light sheet data set before registration (red) and after registration (green). There is swelling ranging from 25-30% with different degrees of swelling along different axes. Note in the sagital image, tissue damage at the top of the cerebellum and substantial physical shift in the brain stem and olfactory bulb. The dedicated LSM to MRH pipeline has been optimized to remove these distortions by bringing the tissue into registration with the original (in the skull) MR volume. The pipeline first down samples the LS to the native resolution of the companion MR data set. Tissue extraction from the skull, even by the most skilled, can result in tissue damage. The human step pairs the MRH and LSM together in an application constructed in Slicer allowing one to place a limited number (<20) fiducials in each volume which drives a label based gross alignment in ANTs. A second step applies a more structured pipeline in ANTS using an initial affine transform followed by a diffeomorphic transform. The resulting transform is then scaled and applied to the full resolution (1.8× 1.8 × 4.0 μm) volumes. Since the three channels are in registration during acquisition, the same transform can be applied to all three full resolution volumes. This operation is executed on the LSM server taking advantage of the 1.5T of RAM. The transform is typically generated in < 3 hrs. Application of the transform to all three volumes can be completed in < 6 hrs. Tiff files are converted to .ims, the proprietary format for Imaris and curated into a directory on the CTX01 (Figure S3) to enable simultaneous visualization of the registered volumes. Figure 4A-C shows a representative axial slice demonstrating registration of NeuN and DWI for specimen 200316-1:1; A) combined NeuN/DWI; B) DWI; C) NeuN. The arrow highlights a region in the edge of the dentate gyrus that is ~ 1 cell wide. The same region is resolved in the DWI image. Figure 4D-F shows the agreement between the radial diffusivity and myelin basic protein (MBP) images. Based on these studies, it became clear that the NeuN LSM data could be used to drive the registration to the DWI MRH images.

### Validation of MRH/TDI vs LS/Thy1

Track density images were generated in DSI studio using a super sampling algorithm described by Calamante et al [22]. The algorithm has been previously validated against conventional histopathology images in a C57BL/6J mouse [23]. Data in these previous studies were acquired with 100 ◻m isotropic spatial resolution, 30 angles, i.e. a resolution index of ~30,000. Using super resolution, Calamante extended the spatial resolution for the TDI to 20 ◻m by seeding the whole brain to 4 million fibers and tracking with a step size of 0.1 mm discarding tracks less the 0.4 mm. The resolution index in specimen 190415-2:1 Table 1 is 108/(.015)^3^ i.e. 18×10^6^, roughly 600 times that of Calamante’s work. Whole brain tractography was acquired in DSI Studio on CTX01 seeding the brain with 54 million fibers. The step size was set deliberately low to .075 mm at a QA threshold of 0.1 percent of the peak QA histogram. Light sheet data were transformed to the MR space using the auto fluorescent image that accompanied the YFP image. The resulting large arrays (TDI~ 4 GB, LS~130GB) were examined in registration in Imaris with representative images shown in Figure 5. To facilitate the registration both TDI and LS are rendered with the same interpolated slice thickness of 14.4 ◻m.

### Validation in a BXD89

Our long-term goal is a methodology to derive a comprehensive collection of IDP in young and old animals across multiple strains of the BXD cohort for experimental system genetics. The final experiment summarized in Table 1 shows two BXD 89 specimens that have been passed through the pipelines described above. Based on the experience with the atlas generation, it became clear that the auto fluorescent channel was not required for LSM to MRH registration. This enabled acquisition of an additional IHC data set. LS data for these specimens were acquired with NeuN, Syto16 and IBAI

## Discussion

Several groups have combined MRI and light sheet imaging. The methods we have described here differ from previous work in four specific ways: 1) the spatial resolution of our MRH data is at least 1000 X higher than that of previous work; 2) the contrast resolution of the MRH data is significantly higher; 3) the MRH data are acquired with the brain in the skull assuring the highest geometric accuracy; 4) the throughput is targeted at neurogenetic studies which require large numbers of specimens. There is still room for improvement. The acquisition time is still too long. The highest resolution 15 ◻m/108 angle volume require ten days, clearly at odds with the need for higher throughput. The reference atlases were acquired with this protocol to assure the highest possible fidelity particularly in generating the connectomes. But recent simulations have suggested that fewer angular samples are required when the spatial resolution is as high as we have achieved. Work is under way to understand tradeoff between spatial and angular resolution, b values, gradient loading, SNR and fidelity of the connectome. Figure S6 shows a comparison between a 25◻m/61 angle protocol and the 15 ◻m atlas (200316-1:1). By reducing the TR to 80 ms and eliminating 2 baseline images, the acquisition time has been reduced to 40 hrs. Work is also underway to implement a magnetization prepared sequence which could provide at least a 2X acceleration. Our first comprehensive connectome study @ 43◻m/121 angles had a resolution index of 1.5×10^6^. The 25 ◻m/61 angle protocol has a resolution index of 3.9×10^6^ with an acquisition time of 40 hrs so our efficiency is improving.

The methods described here rely on a fundamental redesign of our hardware and software infrastructure to enable collection and registration of different data types-anatomic structure from MGRE/DTI, cytoarchitecture from AD/RD/FA, connectomes, regional volumes and cell counts. We believe this approach will offer exciting new ways to understand the genetics that underpin differences in cells and circuits in the brain.

---

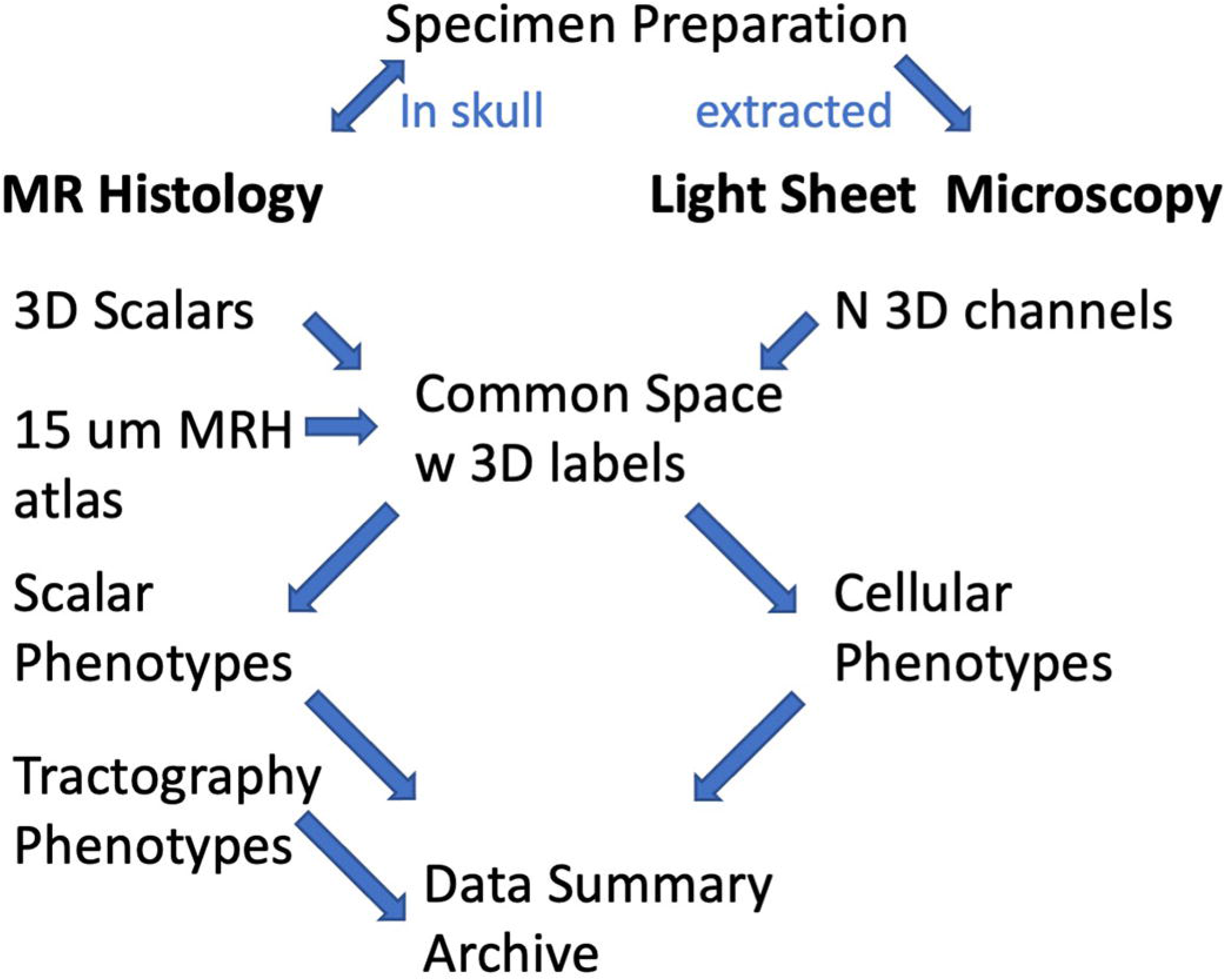

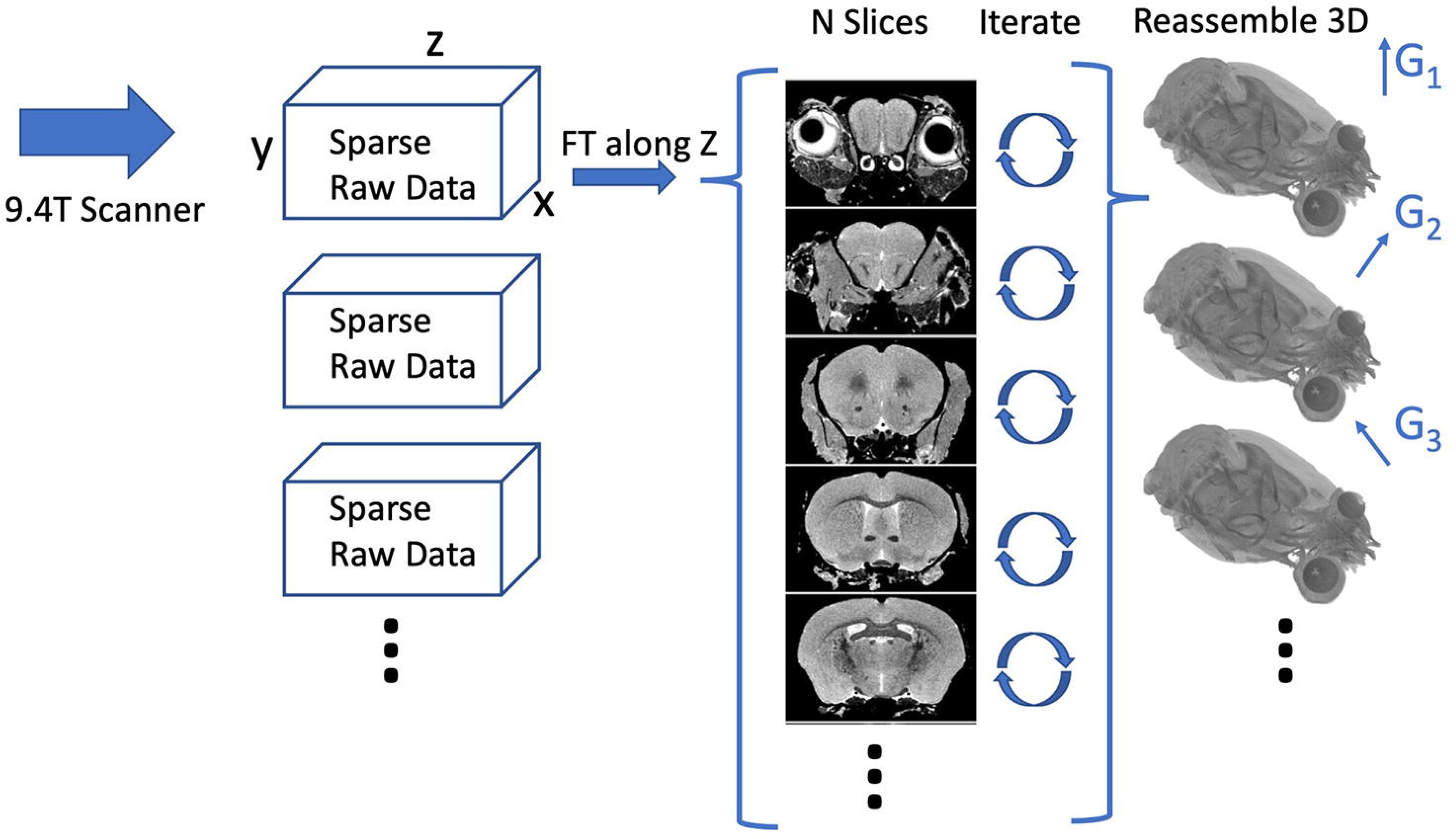

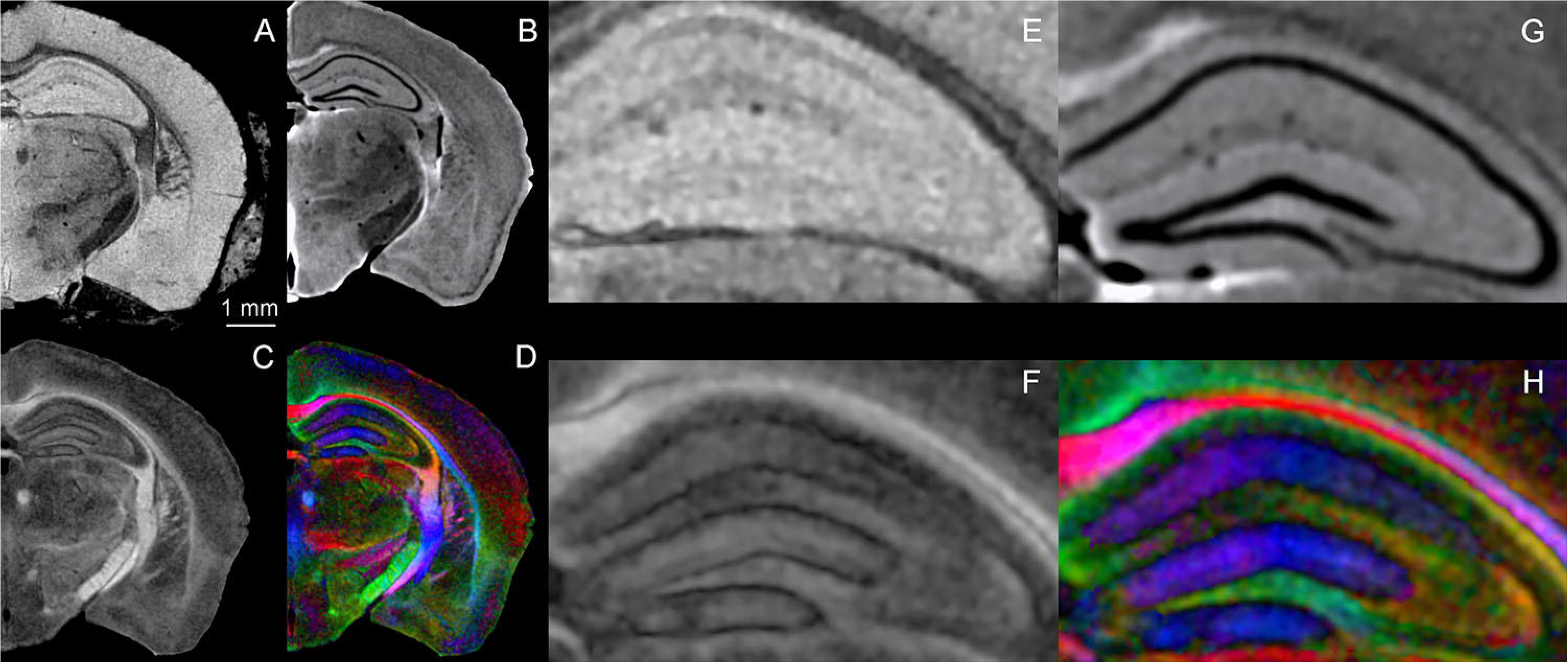

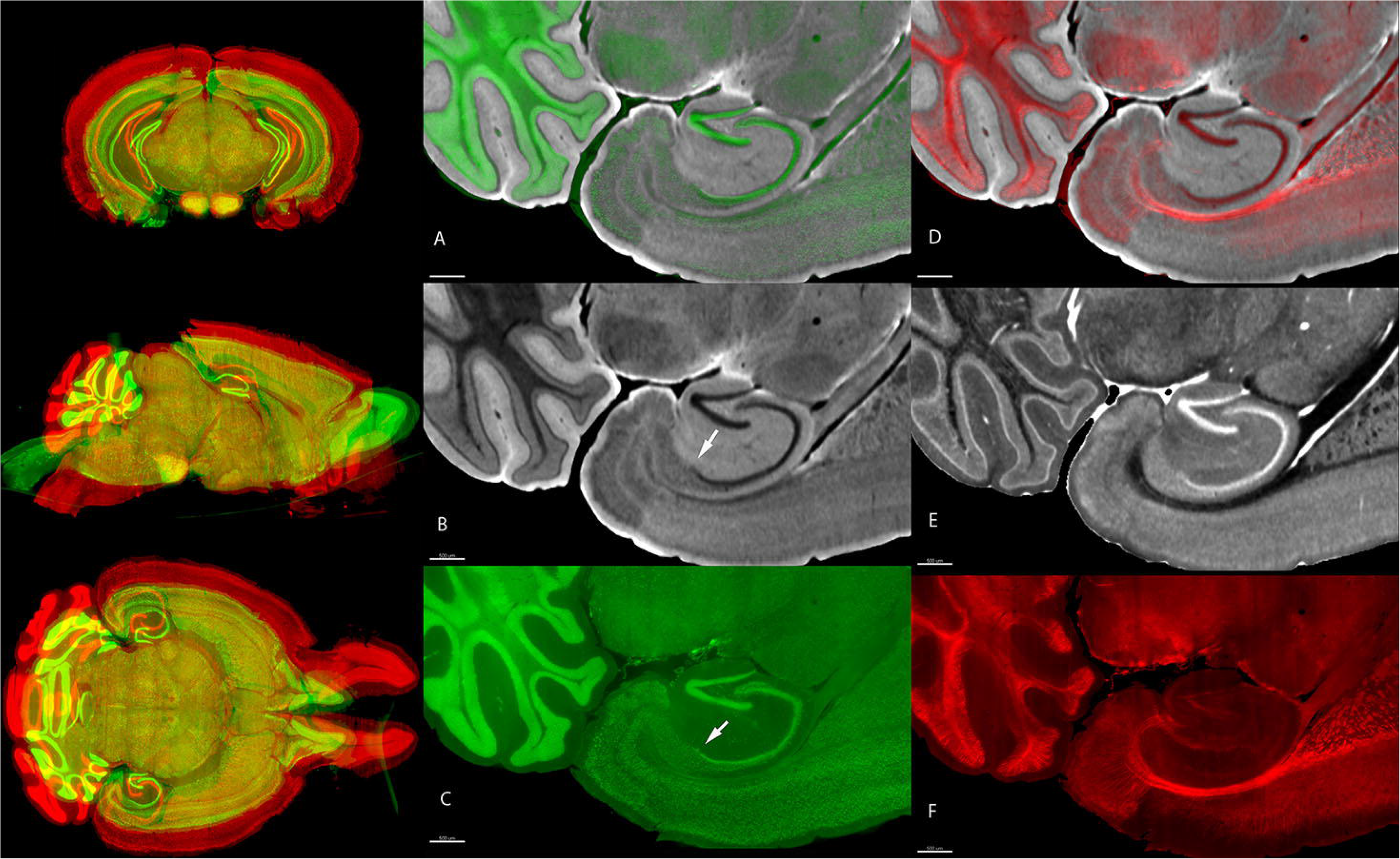

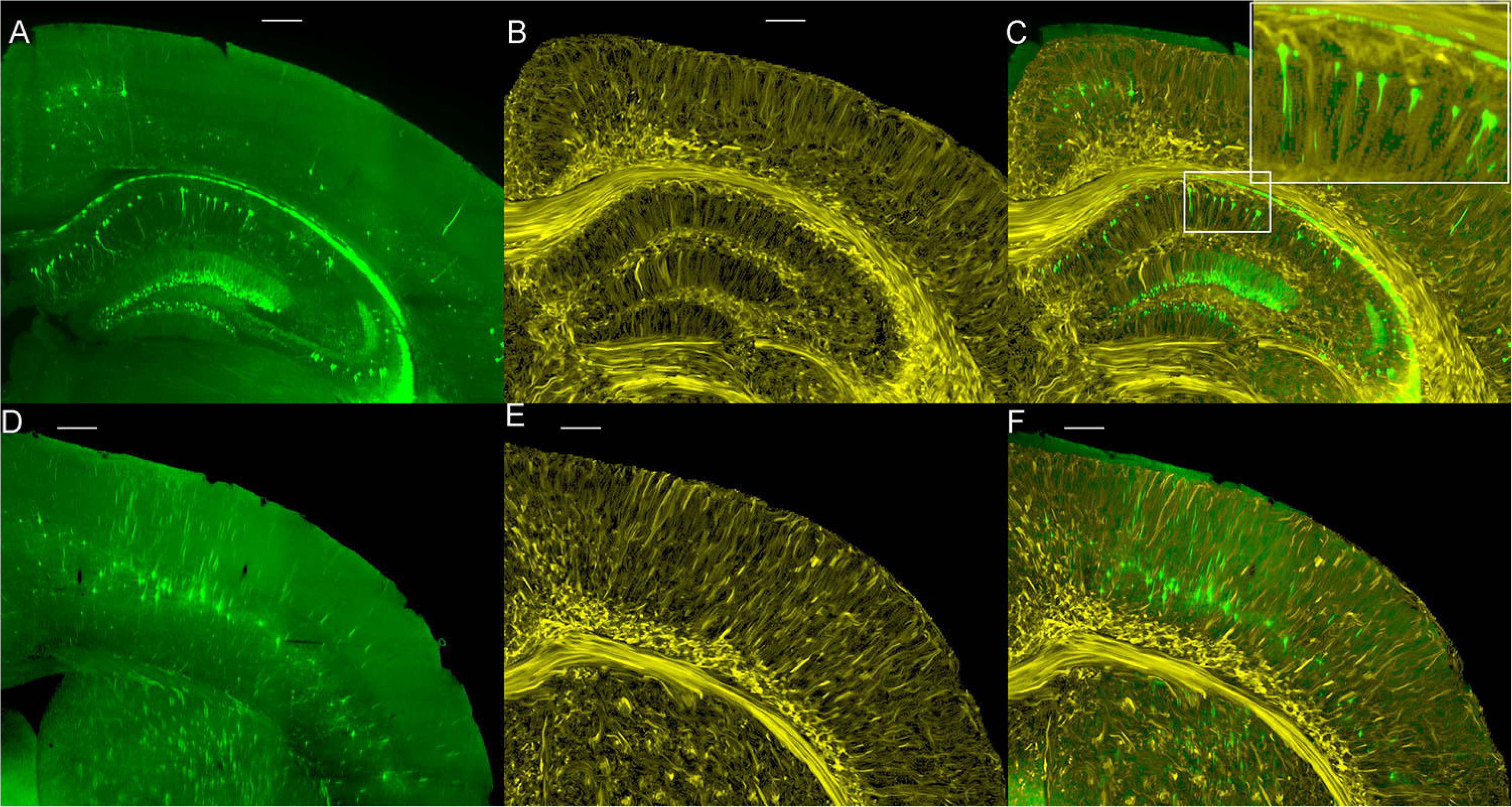

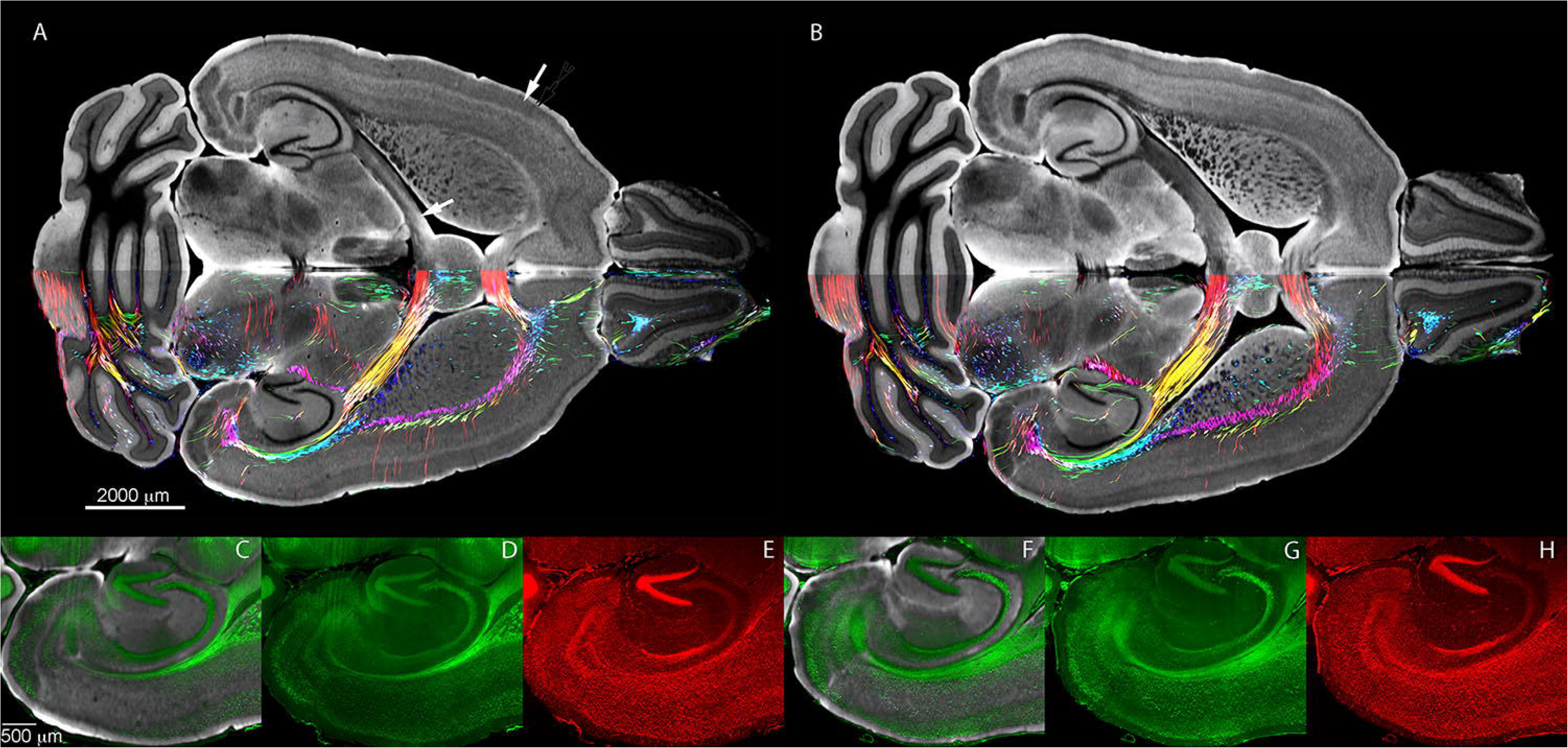

| Resolution Experiments |  | Strain: B6.Cg-Tg (Thy1-YFG)HJrs/J |  |  |  |  |  |
| --- | --- | --- | --- | --- | --- | --- | --- |
| Specimen ID | Gender | Spatial Res (μm) | Angular Res | Time (hrs) | MGRE | Volume (GB) | Light Sheet |
| 190415-3:1 | Male | 25 | 61 | 60 | No | 25 | Thy1-YFG/AutoF |
| 190415-4:1 | Female | 25 | 61 | 60 | No | 25 | Thy1-YFG/AutoF |
| 190415-5:1 | Female | 25 | 61 | 60 | No | 25 | Thy1-YFG/AutoF |
| 190415-1:1 | Male | 25 | 126 | 126 | No | 51 | Thy1-YFG/AutoF |
| 190415-6:1 | Female | 25 | 126 | 126 | No | 51 | Thy1-YFG/AutoF |
| 190415-2:1 | Male | 15 | 61 | 135 | No | 84 | Thy1-YFG/AutoF |
| Atlas Creation |  | Strain: C57BL/6J |  |  |  |  |  |
| 200302-1:1 | Male | 15 | 108 | 245 | Yes | 252 | NeuN/MBP/AutoF |
| 200316-1:1 | Female | 15 | 108 | 245 | Yes | 252 | NeuN/MBP/AutoF |

| Aging Comparison |  | Strain: BXD 89 |  |  |  |  |  |  |
| --- | --- | --- | --- | --- | --- | --- | --- | --- |
| Specimen ID | Gender | Age (days) | Spatial Res (mm) | Angular Res | Time (hrs) | MGRE | Volume (GB) | Light Sheet |
| 190108-5:1 | Male | 111 | 15 | 108 | 245 | Yes | 252 | NeuN/ Syto16/ ibai |
|  |  |  | 45 | 46 | 11.4 | Yes |  |  |
| 200803-12:1 | Male | 687 | 15 | 108 | 245 | Yes | 252 | NeuN /Syto16/ ibai |

