## Supplementary figures and images for "Multimodal Magnetic Resonance Histology and Light Sheet Imaging for Quantitative Neurogenetics of the Mouse"

### Supplemental Figure 1

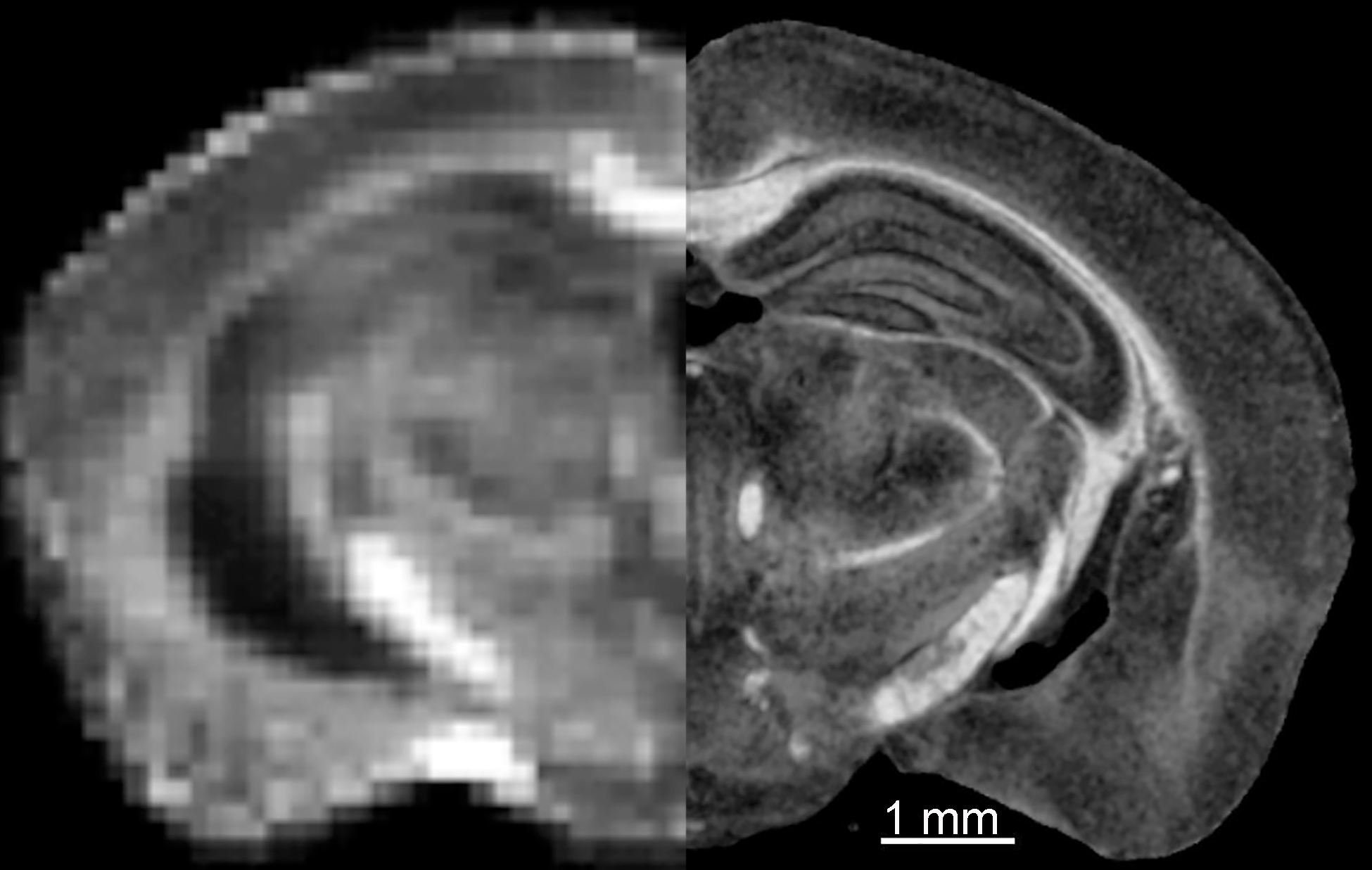

### Supplemental Figure 2

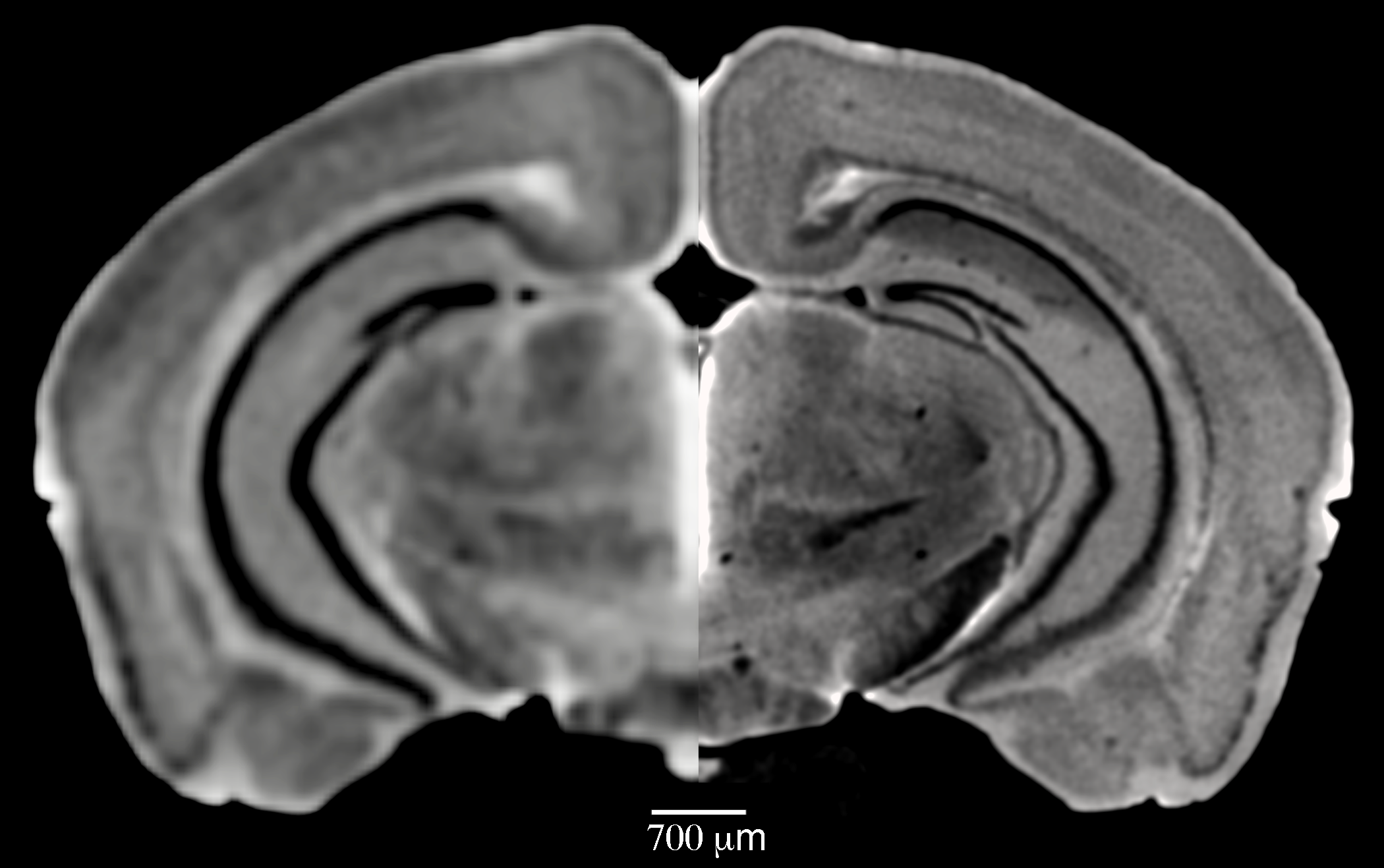

### Supplemental Figure 3

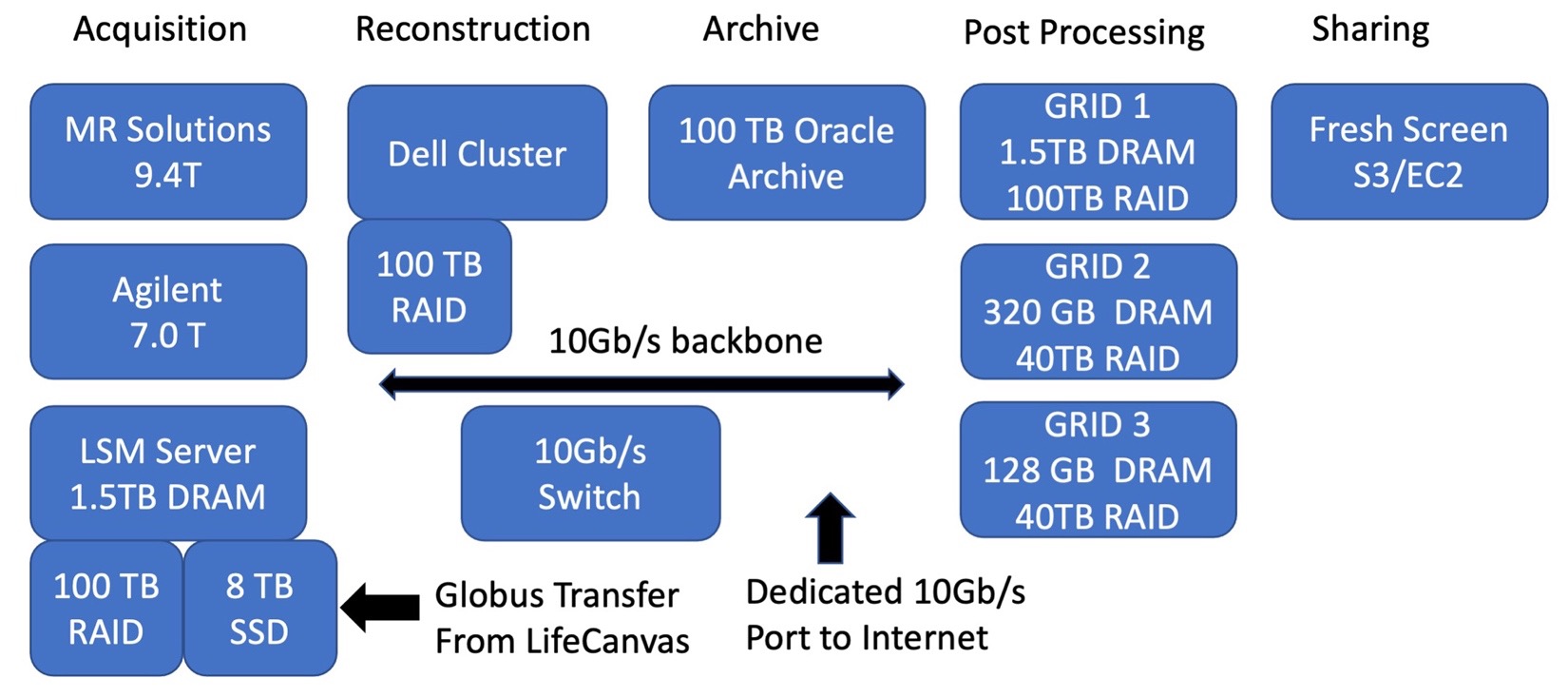

### Supplemental Figure 4

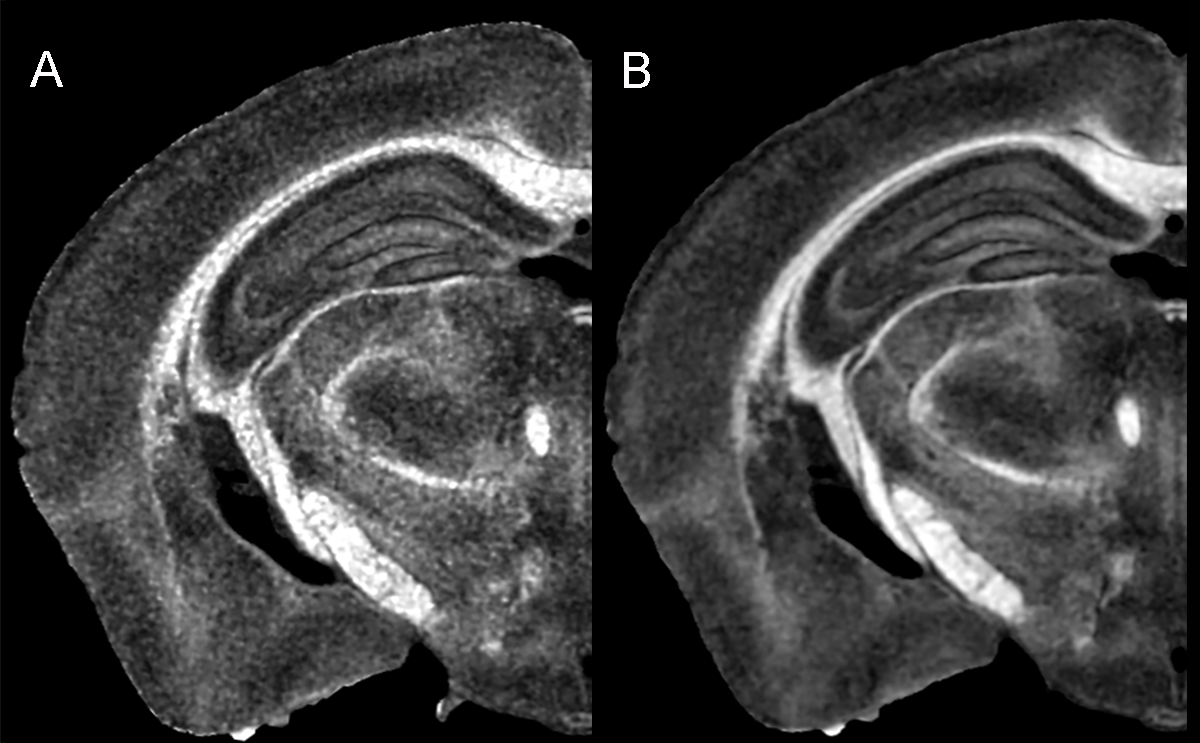

### Supplemental Figure 5

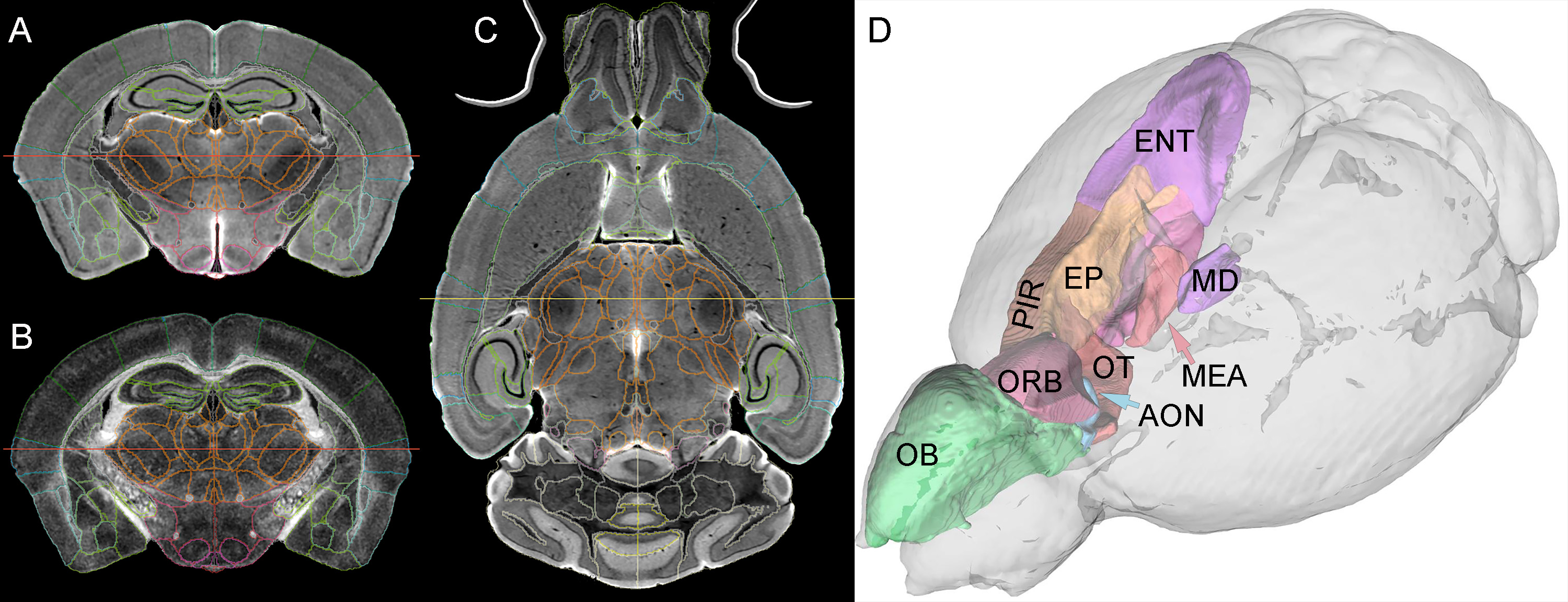

### Supplemental Figure 6

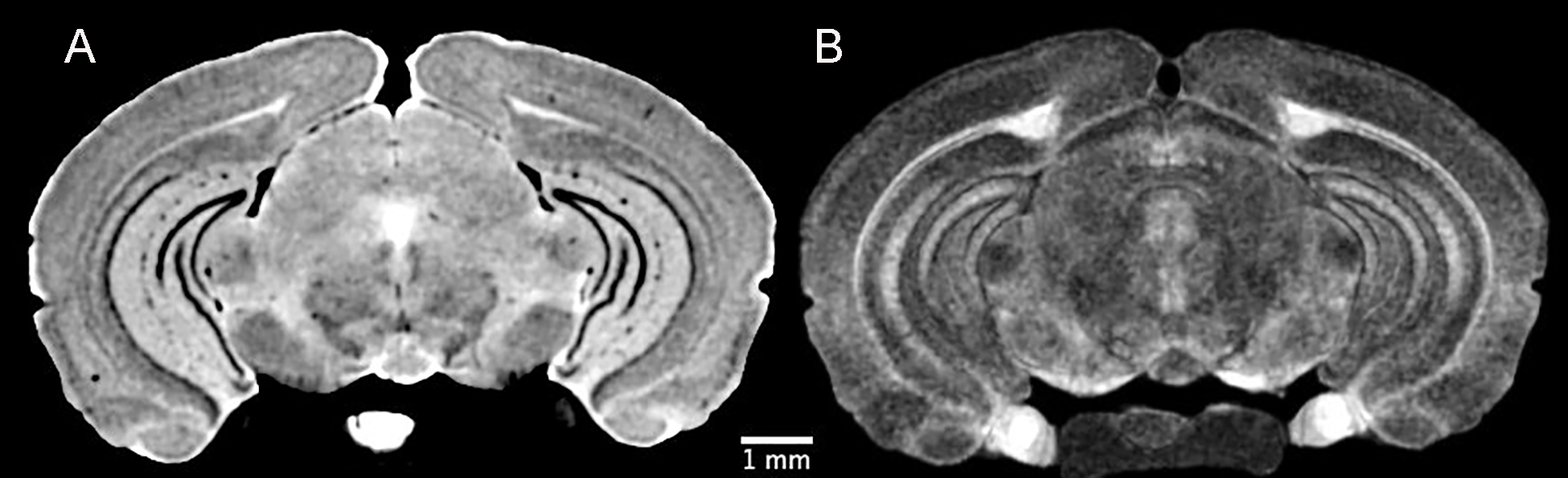
